# The Costs and Benefits of a Modified Biomedical Science Workforce

**DOI:** 10.1101/2020.09.09.289785

**Authors:** Michael D. Schaller

## Abstract

Analyses of the biomedical research workforce, the biomedical research enterprise and its sustainability have identified a number of threats and offered many solutions to alleviate the problems. While, a number of these solutions have been implemented, one solution that has not been broadly adopted, despite being widely recommended, is to increase the number of staff scientists and reduce dependency on trainees. The perceived impediment of this is the cost. This paper explores the costs associated with laboratory personnel and the benefits, in terms of productivity, associated with different positions in the workforce. The results of this cost-benefit analysis depend upon the values assigned to different metrics of productivity by individuals and institutions. If first and senior author publications are the most important metrics of productivity, a trainee dependent workforce is much more cost effective. If total publications is the most valued metric of productivity, the cost effectiveness of trainee and staff scientists is reasonably equitable. This analysis provides data for consideration when making personnel decisions and for the continued discussion of modification of the biomedical research workforce. It also provides insight into the incentives for modification of the workforce at the grass roots, which must be considered by institutions genuinely committed to workforce modification to sustain the biomedical research enterprise.

## INTRODUCTION

Within the past decade, a number of reports raised concerns about the sustainability of the biomedical research enterprise (1–4). These concerns were fueled by the flat NIH budget following its doubling and studies documenting increased numbers of predoctoral and postdoctoral trainees as the number of academic faculty positions remained constant (4–6). The growing workforce coupled with the limits on funding generated a hypercompetitive environment. A consequence of the increased number of trainees qualified for a static number of academic positions was an increase in duration of training period (particularly for postdocs)(4, 7), delaying career advancement and increasing the age of initial funding by an NIH research grant (3, 8, 9).

Consensus recommendations for sustaining biomedical research included increasing sustainable and predictable NIH funding, reducing regulatory burden and modifying training and the workforce (10). Specific recommendations for the workforce included increasing compensation for postdocs, reducing training periods, incorporating professional skills development into training, shifting trainee support to individual and institutional training grants and increasing the use of staff scientists (2, 6, 10–13). There have been notable changes since these recommendations were made. The NIH budget has now increased for five consecutive years with bipartisan and bicameral support in Congress. In response to proposed changes to the Fair Labor Standards Act, the National Research Service Award stipend recommendations were increased, significantly increasing compensation for some postdocs. Increasing awareness of the length of training periods has led to significant efforts to make the metrics of graduate education and postdoctoral training transparent, e.g. the Coalition for Next Generation Life Science initiative, to stimulate identification of mechanisms to reduce the duration of training without compromising quality (14). Numerous reports identify important professional skills that will be useful for trainees regardless of their final choice of profession (12, 15, 16). The NIH Director’s Biomedical Research Workforce Innovation Award: Broadening Experiences in Scientific Training (BEST) supported efforts at 17 institutions to develop innovative approaches to prepare trainees for a broader range of careers (17, 18). Collectively, these efforts have produced changes alleviating some of the issues viewed as threats to the enterprise.

There has been little progress in efforts to support increasing staff scientist positions in the workforce. A staff scientist can be considered a PhD recipient, who has completed postdoctoral training and is working as a full-time employee with benefits performing research. Since 2016, the National Cancer Institute has supported a mechanism providing salary support for extramural staff scientists, however this program is small, with only 84 awards made through the end of FY19. Resistance to this change in the workforce is based upon concerns about the cost and productivity. A priori, one expects a greater cost, but also more productivity, i.e. more papers and citations, with an increase in seniority of the workforce. Obviously, the cost associated with a more senior workforce is due to salary but is also due to benefits available to employees that are unavailable to students and may or may not be available to postdocs. The productivity gain is manifested in an increase in publications and citations. Estimates of the magnitude of the differences between the costs and productivity of individuals at various stages in the biomedical workforce are not widely available. This analysis was undertaken to determine the costs and benefits associated with changing the balance of the biomedical workforce to inform decisions at the individual and institutional level regarding hiring staff scientists.

## MATERIALS AND METHODS

### Salaries and benefits

Public sources were utilized to estimate salaries and benefits of different positions in the biomedical workforce. Sources of salary information included the Ruth L. Kirschstein National Research Service Award (NRSA) stipend levels effective for FY20 (19) and FY18 (20), the Office of Personnel Management (21) and the AAMC Faculty Salary Report (22). Graduate student salaries were also determined by surveying School of Medicine websites for PhD stipend information, yielding stipend levels from 69 institutions, which were averaged to provide an estimated stipend. Benefits available to postdocs were from the National Postdoctoral Association (23) and the costs of benefits at academic institutions were from a survey by the American Association of University Professors (24).

### Publications and citations

Award data and publications affiliated with each award was retrieved from NIH Reporter (https://projectreporter.nih.gov/reporter.cfm). Publication data was curated to remove publications that did not include the PI of the award in the author list and correct misspellings of the PI’s name in the author list to facilitate the analysis. This impacted a small number of publications. Between 1.28% and 2.19% of the publications were culled and 2-3% of the publications required spelling corrections. A python script was used to tally the number of publications associated with each unique award, identify publications on which the PI was first author or senior author and tally each. Publications were pro-rated based upon the duration of each award. Two F31 awards and five awards were excluded since the duration of these awards could not be determined.

Citations were enumerated using a python script to input the PMIDs of the PI’s publications into Pubmed and retrieve the list of PMCIDs of papers that cite the publication. The number of PMCIDs for each publication were tallied. This analysis was performed for first author and senior author publications by each PI.

The distributions of the publication and citation data were not Gaussian and the variance of different samples was not homogeneous. Therefore, nonparametric statistical analyses were performed. For multiple comparisons, the data was analyzed using the Kruskal Wallis H test and Conover and Dunn post hoc tests. The results were considered significant if p<0.01. The analysis was performed using the stats package from SciPy in Python.

The design of this study was reviewed and approved by the West Virginia University Institutional Review Board (WVU Protocol #: 2005003821).

## RESULTS

### Cost – Survey of Salaries and Benefits

The cost associated with changing the demographics of the biomedical workforce is primarily the difference between the salaries and benefits paid to graduate students, postdocs and staff scientists. Salary estimates are provided for each position based upon guidelines from the NIH, surveys of salaries from publicly available data and the Association of American Medical Colleges (AAMC) (Supplemental Table 1). A reasonable estimate of the salary of a biomedical graduate student is $30,305, based upon surveying medical school graduate program web sites. An NIH pay scale for postdocs is based upon experience, with an entry level postdoc earning $52,704 and a postdoc with seven full years of experience earning $64,008 (for FY20) (19). According to the most recent NSF Survey of Doctorate Recipients from US Universities 2018, the median entry salary for biomedical postdocs was $48,000 (25), which was comparable to the FY18 NIH postdoc payscale, where the starting salary was $48,432 (20). Staff scientists at the NIH are paid on the General Schedule system (21). This schedule has 15 grades and 10 steps of salary incrementation per grade (~3%). The entry level for a staff scientist is GS13 and NIH has developed guidelines for elevation of staff scientists to GS14 and GS15. The 2020 compensation for step 1 of GS13 is $78,681 as a base salary and at step 10 is $102,288. In addition to their base salary, most government employees may also receive locality pay to offset discrepancies in pay between the government and the private sector. The Office of Personnel Management of the federal government reports that it takes on average 18 years to advance from step 1 to step 10 in a pay grade (26). There are no data on staff scientist salaries in academia, although there are several sources describing salaries of other positions. The most recent AAMC survey of faculty salaries indicates a median salary of $60,000, and a 75^th^ percentile salary of $72,000 for Instructors (22). According to the American Association of University Professors faculty compensation survey, the average salary of Instructors at doctoral-granting Universities is $65,919. Lecturers’ salaries average $67,896 and faculty with no rank average $79,383 (24). Since staff scientists may currently be categorized under these ranks, these salaries were also considered. The salaries used to measure the cost of different positions is shown in Table 1.

**Table 1.** Estimated costs for labor in the workforce.

| Position | Salary | Benefits | Cost |
| --- | --- | --- | --- |
| Grad Student | \$30,305 | \$3,334* | \$33,639 |
| Grad Student (tuition incl.) | \$30,305 | \$21,750*~ | \$52,005 |
| Postdoc (entry) | \$52,704 | \$11,490*^ | \$64,194 |
| Staff Scientist (entry) | \$78,681 | \$23,353# | \$102,034 |
| *Medical benefit at 11% salary, ~ tuition of \$18,416, ^Retirement benefit at 10.8% salary, #Full benefits at 29.68% | | | |

While reasonable estimates of costs associated with salary can be determined, estimating the costs associated with benefits is more challenging. Typically, graduate student benefits include health insurance. Some institutions include tuition remission when discussing graduate student benefits. The most recent data from the National Center for Education Statistics (for the 2016-17 academic year) indicates that the average in-state graduate tuition was $18,416 (27). At public institutions the average was $11,617 and at non-profit private institutions, the average tuition was $26,551. While the cost of training is real (covered by tuition), this cost is often born by the institution, and may not be part of the calculation of costs by an individual Principal Investigator considering modifying the laboratory workforce. However, this could be a consideration by institutions in evaluating how to invest in the biomedical workforce in future.

Benefits provided to postdocs are extremely variable. The National Postdoctoral Association recently surveyed 190 institutions about issues important to postdocs, including benefits packages available to postdocs (23). The survey parsed postdocs into four categories, postdoc employees (e.g. supported on an R01), postdoc trainees (e.g. supported on a T32), individually funded postdocs (e.g. supported on an F32) and externally funded postdocs (e.g. supported on a foreign fellowship). The benefits available varied greatly between the different categories. Postdoc employees at more than 75% of the institutions were eligible for family health, dental and vision insurance, life insurance, tax-deferred retirement and paid time off. Benefits available to postdocs in other categories are inferior, as illustrated in Supplemental Table 2, which indicates the percentage of institutions providing select benefits to different categories of postdocs (23). The disparity of benefits available to postdocs in different categories is an issue in itself, and illustrates the difficulty in assessing the costs associated with restructuring the workforce. The cost in employee benefits associated with transitioning from a postdoc to a staff scientist could depend upon the source of support for the postdoc. According to data from the National Center for Science and Engineering Statistics, 63.7% of the biology and biomedical sciences postdocs were supported by research grants in 2018 (28), i.e. are classified as postdoc employees by the National Postdoctoral Association. In their 2017 survey, the National Postdoctoral Association found 90% of institutions provided full family medical benefits and 76% of institutions provided tax-deferred retirement benefits to this class of post doc (23). Only 47% of these institutions provided matching retirement benefits to their postdoctoral employees. Therefore, the cost of health insurance and retirement benefits are included in the calculation of the cost of a postdoc.

The estimated costs in benefits for staff scientists was based upon American Association of University Professors faculty compensation surveys (24, 29). Included in these surveys is compensation for instructors and lecturers, academic positions that may be comparable to the staff scientist position. According to the 2019-2020 survey, 95.1% of all full-time faculty at Doctoral Institutions receive medical benefits and 97.1% of faculty receive retirement benefits (24). The average medical benefit provided by the institution is 11.0% of the salary and the average retirement benefit is 10.8%, totaling 21.8% of the salary of the faculty member (Supplemental Table 3). This survey excluded institutional coverage for unfunded retirement liability, prepaid retiree health insurance, social security, long-term disability, Medicare and life insurance. The 2018-2019 survey did not break out individual benefits, but reported salary and total compensation (salary + all benefits). The average benefits across all ranks and all doctoral institutions was 29.68% of the average salary (29). Thus, the results of the 2019-2020 survey underestimate the true cost of benefits. The benefits used in measuring cost for each position is shown in Table 1.

### Benefit - Measurement of Productivity

The benefit associated with changing the demographics of the biomedical workforce is the increase in productivity, on an individual basis, anticipated with the shift to a more experienced workforce. Two measures of productivity were used: 1) publications, specifically total publications, the number of first author publications and the number of senior author publications, and 2) citations, specifically of first author and senior author publications. These metrics readily allow comparison of predoctoral and postdoctoral productivity. Lab and institutional assignments for staff scientists vary, and may or may not include driving a project to first- or senior-author publication. Nevertheless, overall productivity will be captured in the total publications. Identification of cohorts for analysis is challenging, since postdocs have many different titles for the same position (30), a challenge that may also exist in the identification of staff scientists (31). Therefore, the analysis was performed on populations of predoctoral trainees, postdoctoral trainees and staff scientists defined by the NIH, i.e. recipients of individual NIH awards for support of each of the three different positions. A caveat to this strategy is that the analysis may focus upon above average individuals in each category and productivity may exceed the average. In fact, the productivity of F32 awardees is apparently 17% higher than unsuccessful F32 applicants (32).

The National Cancer Institute supports a funding mechanism for staff scientists (PAR-19-291) that began in 2016. The initial comparison was performed on PIs of R50 (staff scientist), F32 (postdoc) and F31 (predoc) awards from the NCI between 2016 and the present. The PIs of K99 (the first stage of Pathway to Independence Awards) awards were also included in this analysis. These cohorts were supported within the same period for cancer-related projects, and thus represented the best match for comparison. The numbers of publications associated with each award, even if published after the end date of the award were captured for this analysis.

The publication profile of the cohorts is shown in Supplemental Table 4. The profiles of the F31 and F32 awardees are similar with ~43% publishing, ~30% publishing a first author paper and only the rare awardee publishing a senior author paper. Approximately 64% of the R50 awardees have published and 9.5% have published a senior author paper. The percentage of R50 awardees publishing a first author paper was similar to the percentage for F31 and F32 awardees (~30%). As a cohort, 71% of the K99 awardees have published a paper, 57% have published a first author paper and 14.4% have published a senior author paper. The statistical analysis of the publication data is summarized in Table 2. Analysis of the total publications revealed no difference in the average numbers of papers per F31 and F32 awardee (0.68 to 1.18 papers per awardee). Total publications by R50 and K99 awardees was significantly different than the F31/F32 awardees but there was no significant difference between the R50 and K99 cohorts (an average of 3.19 to 5.5 total papers per awardee). The F31, F32 and R50 awardees published statistically indistinguishable numbers of first author papers (0.34 to 0.56 papers per awardee), while the K99 awardees published significantly more papers (1.21 papers per awardee). The F31 and F32 cohorts published <0.01 senior author paper per awardee. The R50 and K99 cohorts published more senior author papers (an average of and 0.12 to 0.28 senior author papers per awardee). These results demonstrate increased productivity of staff scientists over predoctorate and postdoctorate trainees in total and senior author publications, but not in first author publications.

**Table 2.** Statistical Comparison of Publication Record of Trainees and Staff Scientists.

| Total Papers Trainees Supported During FY2016 to Present |  |  |  |  |
| --- | --- | --- | --- | --- |
|  | F31s | F32s | K99s | R50 |
| Average number of papers | 0.68 +/- 0.99 | 1.18 +/- 2.05 | 3.19 +/- 5.40 | 5.5 +/- 7.1 |
| 95% confidence interval | 0.58 to 0.78 | 0.86 to 1.49 | 2.30 to 3.99 | 3.97 to 7.03 |
| Kruskal Wallis H test – Statistic = 106.74, p<0.001 |  |  |  |  |
| Dunn post hoc test – F31 = F32, K99 = R50 |  |  |  |  |

| First Author Papers Trainees Supported During FY2016 to Present |  |  |  |  |
| --- | --- | --- | --- | --- |
|  | F31s | F32s | K99s | R50 |
| Average number of papers | 0.34 +/- 0.60 | 0.54 +/- 1.02 | 1.21 +/- 1.98 | 0.56 +/- 1.14 |
| 95% confidence interval | 0.28 to 0.41 | 0.39 to 0.70 | 0.92 to 1.50 | 0.31 to 0.81 |
| Kruskal Wallis H test – Statistic = 55.68, p<0.001 |  |  |  |  |
| Dunn post hoc test – F31 = F32 = R50, K99 is different |  |  |  |  |

| Senior Author<br>Papers Trainees Supported During FY2016 to Present |  |  |  |  |
| --- | --- | --- | --- | --- |
|  | F31s | F32s | K99s | R50 |
| Average number of papers | 0.003 +/- 0.053 | 0.006 +/- 0.078 | 0.28 +/- 1.30 | 0.12 +/- 0.39 |
| 95% confidence interval | -0.003 to 0.008 | -0.006 to 0.18 | 0.087 to 0.47 | 0.035 to 0.20 |
| Kruskal Wallis H test – Statistic = 65.128, p<0.001 |  |  |  |  |
| Dunn post hoc test – F31 = F32, K99 = R50 |  |  |  |  |

The parameters of this comparison were driven by the timing of the R50 Awards to staff scientists. The limitations include the small number of awardees analyzed (84 R50 awardees) and the time frame, which runs to the present. The publications will be under representative as studies in new and ongoing projects are incomplete and not yet published. Citations were not analyzed in these cohorts, given the limited time period since initiation of the awards and the concern that they would grossly under estimate the impact of these publications.

To address these limitations, the analysis was expanded to all NIH F31 and F32 recipients from FY2012 through FY2016. As there was no staff scientist funding mechanism during this time, the analysis was extended to K99 recipients, to measure productivity of a population more advanced than F32 recipients. The analysis of R50 and K99 recipients from NCI from FY16 to the present suggests total publication and senior publications are similar between these cohorts and thus K99 data is a suitable proxy for the analysis. This increased the number of awardees analyzed (8,445 total awardees) and the interval since the time-frame of support provided time for project completion and publication, increasing confidence in the measurement. This also allowed the incorporation of citations into the analysis. There is a small overlap in trainees between support mechanisms. Of the 3605 F31 trainees, 3.1% were also supported by an F32 fellowship during the time period, and 4.1% of the F32 recipients also received K99 awards that were active between the beginning of FY2012 and the end of FY2016.

The publication profiles of these trainees are illustrated in Supplemental Table 5. Obviously, shifting the time frame for analysis back led to a more robust publication profile. Approximately 83% of F31/F32 and 93% of K99 trainees published at least one paper. First author papers were published by 73-75% of F31 and F32 awardees and 84% of the K99 trainees. The percentage F31 and F32 trainees publishing senior author papers was small (2-6%), whereas 34.7% of K99 awardees published senior author papers. Preliminary analysis of an earlier cohort of trainees (F31 and F32 awardees from FY2007 to FY2011) revealed similar publication profiles, suggesting further adjustment of the time frame of study would not yield additional insight. Statistical analysis of the FY2012 to FY2016 cohorts is shown in Table 3. There were significant differences in the total number of papers published by the different cohorts (F31 = 2.66 papers/awardee, F32 = 3.03 papers/awardee, K99 = 4.92 papers/awardee). F31 and F32 awardees published similar numbers of first author papers (1.66 and 1.71 papers/trainee respectively), while K99 trainees published significantly more first author papers (2.01 papers/awardee). Senior author publications differed significantly between F31 (0.03 papers/trainee), F32 (0.08 papers/trainee) and K99 (0.71 papers/trainee) recipients. To normalize for the differences in duration of each fellowship, the productivity of each cohort was pro-rated to generate an average number of papers per year (Figure 1, Table 4). In terms of total papers, the F31 cohort published an average of 1.37 papers per year, F32 trainees published 1.57 papers per year and the K99 awardees published 3.02 papers per year. On average, F31 and F32 trainees published 0.88-0.89 first author papers per year and K99 trainees published 1.21 papers per year. The results demonstrate the expected increase in productivity with academic advancement. There are several caveats to the analysis. The results may overestimate productivity, since the cohorts contain the most highly competitive individuals at each training level. Conversely, the results may underrepresent productivity, since publications are restricted to those associated with the funded project and may be only a partial publication record of any given individual. Data for staff scientists is unavailable for this analysis. By extrapolation from Table 2, the first author publication record of K99 recipients exceeds staff scientists, but the total publications and senior author publications are comparable.

**Figure 1.**
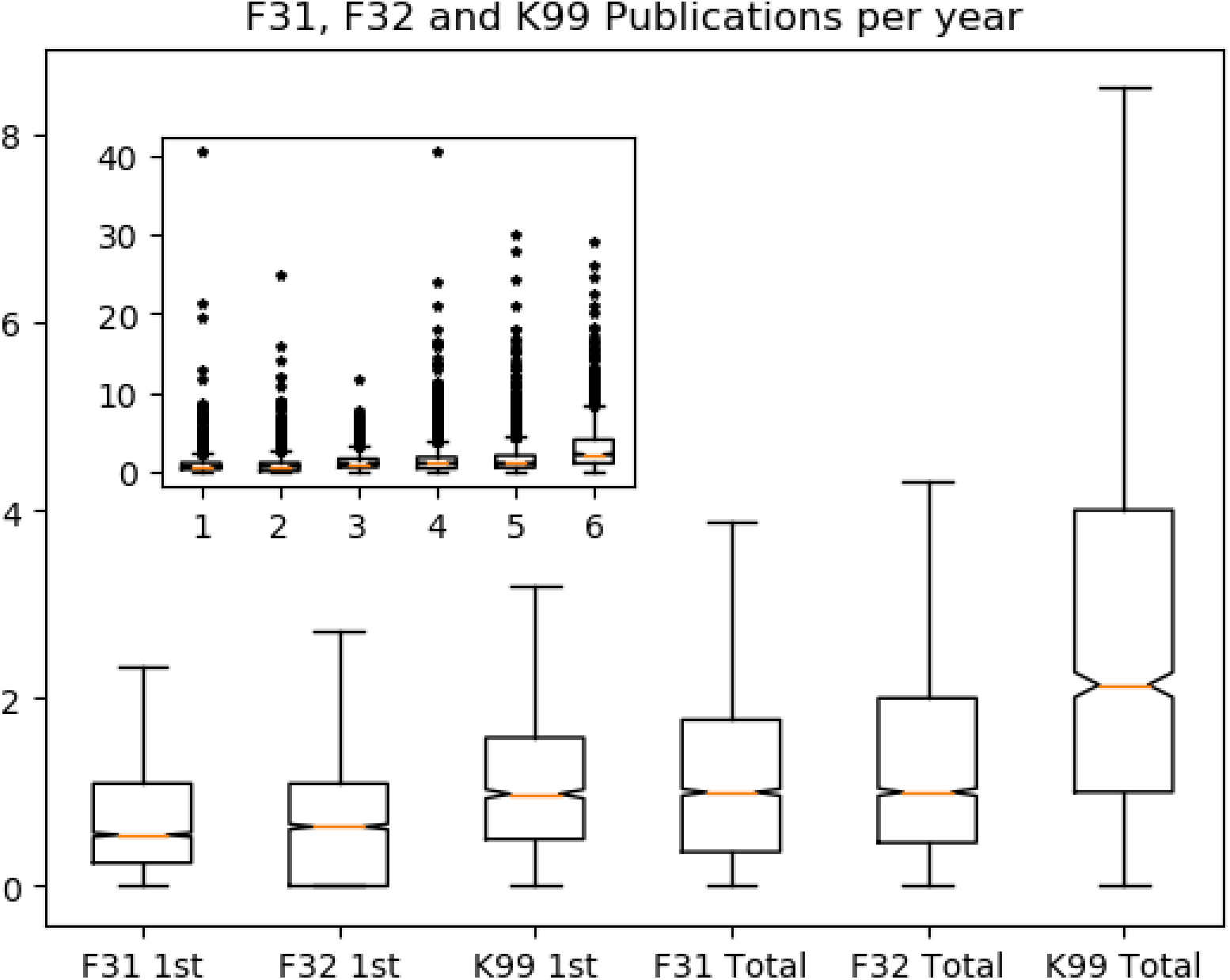
Number of publications per year. The number of 1^st^ author and total publications per year for F31, F32 and K99 awardees is shown in a box and whisker plot. The inset includes the outliers. The number of 1^st^ author publications for K99 awardees is significantly different than the others. The total number of publications is significantly different between each of the groups.

**Table 3.** Statistical Comparison of Publication Record of Trainees.

| **Total Papers Trainees Supported During FY2012 to FY2016** |  |  |  |
| --- | --- | --- | --- |
|  | **F31s** | **F32s** | **K99s** |
| Average number of papers | 2.66 +/- 2.94 | 3.03 +/- 3.28 | 4.92 +/- 4.53 |
| 95% confidence interval | 2.56 to 2.75 | 2.93 to 3.14 | 4.67 to 5.18 |
| Kruskal Wallis H test – Statistic = 375.980, p< 0.001 |  |  |  |
| Dunn post hoc test – F31, F32 and K99 are all different |  |  |  |

| **First Author Papers Trainees Supported During FY2012 to FY2016** |  |  |  |
| --- | --- | --- | --- |
|  | **F31s** | **F32s** | **K99s** |
| Average number of papers | 1.66 +/- 1.83 | 1.71 +/- 1.84 | 2.01 +/- 1.86 |
| 95% confidence interval | 1.60 to 1.72 | 1.65 to 1.77 | 1.91 to 2.12 |
| Kruskal Wallis H test – Statistic = 53.696, p< 0.001 |  |  |  |
| Dunn post hoc test – F31 = F32 and K99 is different from both F31 and F32 |  |  |  |

| **Senior Author**<br>**Papers Trainees Supported During FY2012 to FY2016** |  |  |  |
| --- | --- | --- | --- |
|  | **F31s** | **F32s** | **K99s** |
| Average number of papers | 0.03 +/- 0.32 | 0.08 +/- 0.38 | 0.71 +/- 1.41 |
| 95% confidence interval | 0.05 to 0.07 | 0.09 to 0.12 | 0.74 to 0.92 |
| Kruskal Wallis H test – Statistic = 1287.3, p< 0.001 |  |  |  |
| Dunn post hoc test – F31, F32 and K99 are all different |  |  |  |

**Table 4.** Statistical Comparison of Publication Record of Trainees – Prorated for Length of Award.

| **Total Papers per year - Trainees Supported During FY2012 to FY2016** |  |  |  |
| --- | --- | --- | --- |
|  | **F31s** | **F32s** | **K99s** |
| Average number of papers | 1.37 +/- 1.83 | 1.57 +/- 2.00 | 3.02 +/- 3.15 |
| 95% confidence interval | 1.31 to 1.43 | 1.50 to 1.64 | 2.85 to 3.20 |
| Kruskal Wallis H test – Statistic = 559.038, p<0.001 |  |  |  |
| Dunn post hoc test – F31 is different from F32 is different from K99 |  |  |  |

| **First Author Papers per year Trainees Supported During FY2012 to FY2016** |  |  |  |
| --- | --- | --- | --- |
|  | **F31s** | **F32s** | **K99s** |
| Average number of papers | 0.88 +/- 1.30 | 0.89 +/- 1.22 | 1.21 +/- 1.22 |
| 95% confidence interval | 0.83 to 0.92 | 0.85 +/- 0.93 | 1.14 to 1.28 |
| Kruskal Wallis H test – Statistic = 155.258, p<0.001 |  |  |  |
| Dunn post hoc test – F31 and F32 are the same, K99 is different |  |  |  |

| **Senior Author Papers per year - Trainees Supported During FY2012 to FY2016** |  |  |  |
| --- | --- | --- | --- |
|  | **F31s** | **F32s** | **K99s** |
| Average number of papers | 0.017 +/- 0.17 | 0.04 +/- 0.23 | 0.46 +/- 0.96 |
| 95% confidence interval | 0.01 to 0.02 | 0.03 to 0.05 | 0.41 to 0.51 |
| Kruskal Wallis H test – Statistic = 1316.951, p<0.001 |  |  |  |
| Dunn post hoc test – F31 is different from F32 is different from K99 |  |  |  |

The number of citations for each first author and senior author paper by each trainee was tallied. Citations of first author papers increased with advances in training (Table 5). F31 publications were cited an average of 15.34 times, F32 trainee’s papers were cited an average of 22.07 times and K99 first author papers were cited an average of 26.55 times. These differences are significant. Conversely, citations for senior author publications did not significantly differ between the cohorts. The highest number of citations per paper was for papers by F32 trainees (17.09 citations per paper) and the lowest was for papers published by K99 trainees (12.20 citations per paper). The observation that senior author publication by F32 and K99 trainees are cited less frequently than first author publications is intriguing. The reason for this disparity is unclear.

**Table 5.**
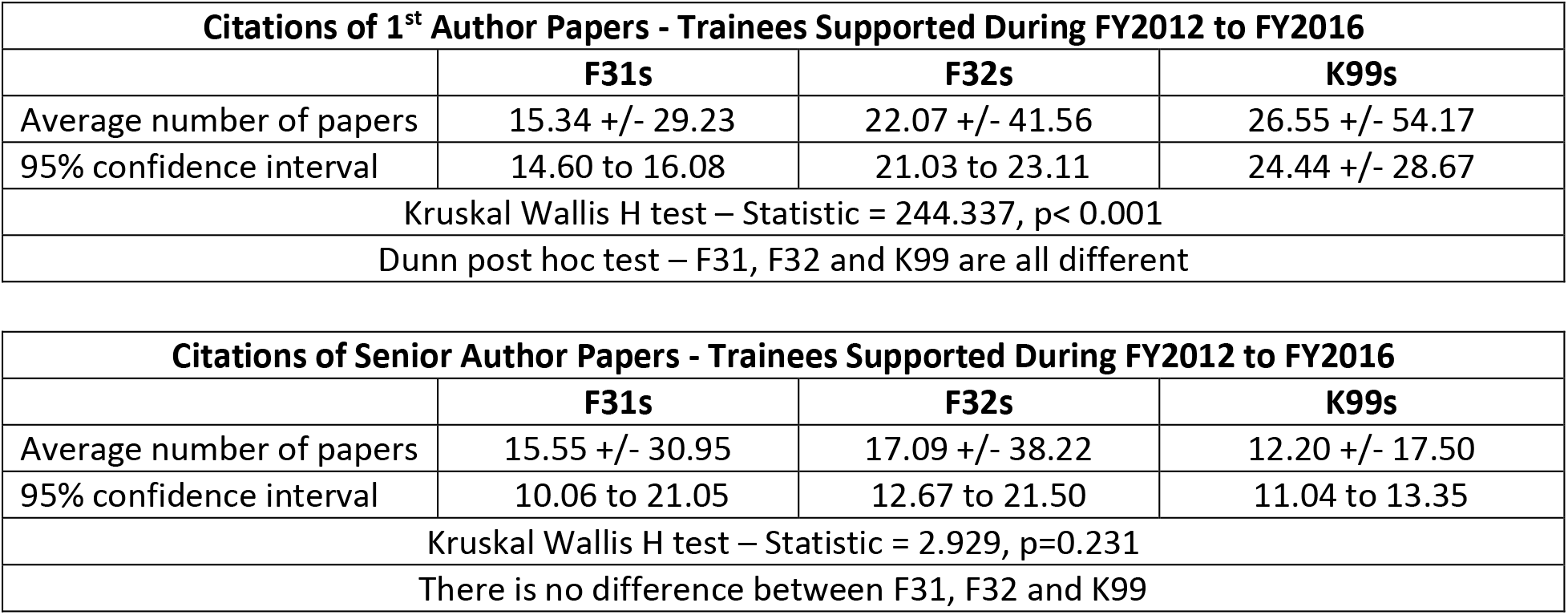
Statistical Comparison of Citation Record of Trainees.

| **Citations of 1<sup>st</sup> Author Papers - Trainees Supported During FY2012 to FY2016** |  |  |  |
| --- | --- | --- | --- |
|  | **F31s** | **F32s** | **K99s** |
| Average number of papers | 15.34 +/- 29.23 | 22.07 +/- 41.56 | 26.55 +/- 54.17 |
| 95% confidence interval | 14.60 to 16.08 | 21.03 to 23.11 | 24.44 +/- 28.67 |
| Kruskal Wallis H test – Statistic = 244.337, p< 0.001 |  |  |  |
| Dunn post hoc test – F31, F32 and K99 are all different |  |  |  |

| **Citations of Senior Author Papers - Trainees Supported During FY2012 to FY2016** |  |  |  |
| --- | --- | --- | --- |
|  | **F31s** | **F32s** | **K99s** |
| Average number of papers | 15.55 +/- 30.95 | 17.09 +/- 38.22 | 12.20 +/- 17.50 |
| 95% confidence interval | 10.06 to 21.05 | 12.67 to 21.50 | 11.04 to 13.35 |
| Kruskal Wallis H test – Statistic = 2.929, p=0.231 |  |  |  |
| There is no difference between F31, F32 and K99 |  |  |  |

### Costs versus Benefits

The costs and benefits associated with a lab workforce comprised of different combinations of predoctoral and postdoctoral trainees or a staff scientist are presented in Table 6. The estimated total compensation for an entry level staff scientist on the G13 pay scale and an entry level postdoc are included. The estimated total compensation for a predoctoral trainee including and excluding tuition is shown. Financial planning at the lab level most often does not include a consideration of tuition, whereas tuition is a component of institutional planning. The compensation of an entry level staff scientist is equivalent to the compensation of 3.02 graduate students, excluding tuition, 1.95 graduate students, including tuition, and 1.58 postdoctoral trainees. Using entry level data minimizes the cost differential. For comparison, the compensation of a staff scientist at step 6 of the G13 pay scale is equivalent to the compensation of 3.54 graduate students, excluding tuition, 2.29 graduate students, including tuition, and 1.85 postdoctoral trainees.

**Table 6.** Cost-Benefit Analysis.

| **Costs associated with workforce** |  |  |  |  |  |  |
| --- | --- | --- | --- | --- | --- | --- |
|  | **1 staff scientist** | **2 PhDs** | **3 PhDs** | **1 PDF** | **2 PDF** | **1 PhD + 1 PDF** |
| **Salary + Benefits** | \$101,439 | \$67,278 | \$100,917 | \$64,194 | \$128,388 | \$97,833 |
| **Salary + Benefits + Tuition** | \$101,439 | \$104,110 | \$156,165 | \$64,194 | \$128,388 | \$116,249 |
| **Benefits associated with the workforce** |  |  |  |  |  |  |
|  | **1 staff scientist** | **2 PhDs** | **3 PhDs** | **1 PDF** | **2 PDF** | **1 PhD + 1 PDF** |
| **Total papers** | 3.02 | 2.74 | 4.11 | 1.57 | 3.14 | 2.94 |
| **1<sup>st</sup> Author** | 0.89* | 1.76 | 2.64 | 0.89 | 1.78 | 1.77 |
| **Sen. Auth.** | 0.46 | 0.034 | 0.051 | 0.04 | 0.08 | 0.097 |
| **Citations for 1<sup>st</sup> Author** | 28.59 | 30.68 | 46.02 | 22.07 | 44.14 | 37.41 |
| **Citations for Sen. Auth.** | 19.98 | 31.1 | 46.65 | 17.09 | 34.18 | 32.64 |
| **Total Cit.** | 48.56 | 61.78 | 92.67 | 39.16 | 78.32 | 70.05 |
| The costs are derived from salaries, benefits and tuition presented in Table 1. The benefits are derived from the pro-rated publications presented in Table 4 and citations presented in Table 5. The publications/citations for K99 awardees are used as a proxy for staff scientists. * since the number of 1<sup>st</sup> author papers by staff scientists appears statistically different than 1<sup>st</sup> author papers by K99 awardees, but comparable to postdoctoral trainees, the number of 1<sup>st</sup> author papers by postdocs is used. |  |  |  |  |  |  |

The data for benefits associated with each position in Table 6 are derived from Tables 4 and 5. Data for K99 awardees is used as a proxy for staff scientists based upon the similarity in productivity, shown in Table 2, with the exception of 1^st^ author publications. As the number of first author publications of staff scientists is not statistically different from the number of 1^st^ author publications of trainees (Table 2), the number of 1^st^ author publications for postdocs (Table 4) is used for staff scientists in Table 6. The total benefits of these combinations of scientists is composed of four components, total papers, 1^st^ author papers, senior author papers and total citations. Comparison of total publications per year shows that the productivity of a staff scientist is comparable to the productivity of the combination of 2 predoctoral trainees or two postdoctoral trainees, or a predoctoral and postdoctoral trainee. The total publications per year of 3 predoctoral students exceeds that of a staff scientist. From this perspective the cost differential is similar to the benefit differential. The number of 1^st^ author publications per year of any combination of more than one predoctoral or postdoctoral trainee, exceeds the number of 1^st^ author publications of a staff scientist. In all of the workforce scenarios presented, the staff scientist publishes more senior author papers. If primary publications, i.e. the sum of 1^st^ author and senior author publications, are considered, any combination of more than one predoctoral or postdoctoral trainee publishes more primary publications than a staff scientist. From this perspective, the costs of a staff scientist exceed the benefits. Finally, the number of citations per year of any combination of predoctoral and postdoctoral trainees exceeds those of a staff scientist. This is a derivative of the number of 1^st^ author publications produced by each group.

## DISCUSSION

This report compares salaries and estimated total compensation between different positions in the biomedical workforce to evaluate the costs associated with modifying the workforce. While the comparison was made using entry level positions and minimizes the cost differential, data within the report and source material allow similar comparisons of other scenarios. Estimates of the benefits associated with modifying the workforce are based upon productivity, i.e. publications and citations. These estimates are based upon the records of recipients of individual NIH training or staff scientist awards and may be reflective of above average individuals. The interpretation of this analysis depends upon the value associated with 1^st^/senior author publications and middle author publications. The contribution of 1^st^ and senior authors to publications is apparent, and if this metric is used as the benefit, moving from trainees to staff scientists is a losing proposition. However, if the major consideration is overall productivity, the costs and benefits of a staff scientist are relatively equitable to different combinations of trainees, with the exception of the combination of 3 predoctoral trainees. How the different measures of productivity are weighed by individual principal investigators and institutions will drive their interpretation of the costs and benefits associated with changing the workforce and will inform their decisions.

This comparison of productivity indicates that doctoral trainees may contribute on average 1.37 papers per year, postdoctoral trainees a total of 1.57 papers per year and staff scientists (using K99 awardees as surrogates) a total of 3.02 papers per year. There are several other studies measuring productivity of trainees. A study of 933 science and engineering students at the California Institute of Technology from 2000 to 2012 measured the number of publications per year from 5 years prior to the year of dissertation defense to 2 years after the defense. Peak productivity was in the year prior to the defense and year of defense where the average number of papers was approximately 1.1 (for women) and 1.25 (for men) (33). Their longitudinal analysis illustrated another important consideration, that the productivity of students is initially very low, rises to a peak and then continues after graduation as work initiated by the student is completed and published. The productivity of a long-time staff scientist is expected to be continuous and not subject to the 5- or 6-year cycle of ramping up productivity. A different approach was used to study the contributions of lab members to overall lab productivity from 1966 to 2000 in the Massachusetts Institute of Technology Department of Biology (34). Comparing annual lab productivity to lab composition, the authors suggest that one doctoral student contributed to an increase of 0.14 publications per year and one postdoc contributed 0.34 publications per year. The addition of a research technician to a lab had no impact on the number of publications. The authors extrapolate this observation to indicate that the addition of staff scientists would have little overall impact on lab productivity. However, the argument can be made that research technicians and staff scientists fulfill very different roles and have very different skill sets. Interestingly, research technicians did have an impact on the number of “breakthrough” publications, i.e. papers appearing in Science, Cell or Nature. There have been several studies measuring postdoc productivity. To assess the impact of doubling of the NIH budget upon productivity, a total of 3669 postdocs identified from the Survey of Doctorate Recipients from 1995, 2001 and 2003, were studied (35). Productivity was measured by total number of papers published in the preceding 5 years. The doubling had no effect upon productivity, which averaged 1.08 papers per year. In a study to assess the effect of extramural support upon research productivity, successful applicants for F32 awards between 1980 and 2000 were compared with unsuccessful applicants (32). Productivity was measured as the number of publications in the 5 years following submission of the application. Unsuccessful applicants published 0.92 papers per year, while successful applicants published 1.08 papers per year. Comparison with results in the literature suggests that 1.37 predoctoral papers per year and 1.57 postdoc papers per year might overestimate trainee productivity and thus underestimate the differential in productivity of a staff scientist.

There are three strategies to employ staff scientists in the biomedical workforce. The expectations of staff scientists vary depending upon their assignment and this could impact metrics of productivity. First, the staff scientist could be a position in an individual research lab, providing intellectual capacity, experience and a technical skill set devoted to driving the research program of the lab forward. The lab bears the cost and reaps the benefits of the staff scientist, which could include 1^st^ and senior author publications exceeding the estimates in Table 6. Second, the staff scientist could provide specific expertise and technical skills to the research community, e.g. as the director of a core facility. The costs are distributed broadly across users of the facility and the institution and the users of the facility receive the benefit of the staff scientist’s expertise, which would primarily be middle author publications in collaboration with different labs. The benefits are tangible but may make a relatively small contribution to any single research program. Third, the staff scientist position could be shared among several laboratories, each contributing to the cost and receiving the benefits of the position. A staff scientist in this position could be critical for team science approaches to problems and a key player in Program Project Grants, CoBRE Grants in IDeA states and other developing teams. Supporting staff scientists in these positions would foster the development of team science.

This analysis was undertaken to examine the costs at the individual level associated with changing the workforce to gauge the incentives/disincentives associated with replacing trainees with staff scientists. However, there are other considerations for the scientific community to contemplate. First, the recommendation to increase staff scientists in the workforce was made in concert with other recommendations to modify training, which will impact the workforce. The workforce in the biological and biomedical sciences in academia in 2018 was comprised of 52,627 predoctoral trainees, 21,533 postdoctoral trainees and 8,250 nonfaculty researchers (ref). Some non-tenure track faculty are also part of the workforce. Since the vast majority of the workforce is comprised of trainees, the enterprise continuously produces an excess of skilled researchers. There are jobs for these researchers in academia, industry and government, but approximately 47% of trainees pursue jobs in non-research intensive positions (36). While this analysis shows that the productivity of 3 predoctoral trainees exceeds that of a staff scientist for the same financial cost, the unseen cost to the enterprise is the production of 3 additional skilled researchers to advance in the workforce. Second, increasing staff scientists will increase stability of the workforce. In 2018, 25% of the predoctoral trainees and 54.6% of the postdoctoral trainees in the biological and biomedical sciences were temporary visa holders. The diversity of background of temporary visa holders and their productivity are assets to the biomedical workforce. Forces external to biomedical research can impact this component of the workforce, e.g. policies intended to curb foreign influence in science. Shifting the workforce to support more staff scientists could negate this type of threat to biomedical science.

If primary publications are the valued metric in making these decisions at the lab level, the costs associated with replacing trainees with a staff scientist exceed the anticipated benefits in increased productivity, thus providing no incentive for this transition. Other incentives, e.g. lack of access to graduate students, could drive the incorporation of staff scientists into research programs out of necessity rather than design to reshape the workforce. If there are institutional commitments to redress the workforce imbalance in the biomedical research enterprise, researchers will require incentives to transition the workforce to meet future needs. The National Cancer Institute took the initiative by establishing a support mechanism for staff scientists. Additional Institutes at the NIH should consider implementing similar programs. NIH should also consider supplements to partially defray the costs associated with employing staff scientists on R01 grants (e.g. providing half of the cost) and mechanisms to support staff scientists who are critical to team science initiatives. Academic and research institutions must also play a critical role in any modification of the biomedical workforce. Institutional investments to support staff scientists to promote team science and protect their investment in research against erosion of graduate and postdoctoral programs in response to changes in the research enterprise. Forward thinking and proactive institutions will position themselves advantageously to meet future challenges to the biomedical research enterprise.

## Abbreviations

NIH: National Institutes of Health
BEST: Broadening Experiences in Scientific Training
FY: fiscal year
NRSA: National Research Service Award
AAMC: Association of American Medical Colleges
PMID: PubMed reference number
PMCID: PubMed Central identifier
PI: Principal Investigator
NCI: National Cancer Institute
CoBRE: Centers of Biomedical Research Excellence
IDeA: Institutional Development Award

## ACKNOWLEDGEMENTS

Many thanks to Chris Pickett and Ben Corb for insightful comments and suggestions to improve the manuscript.

## AUTHOR CONTRIBUTIONS

MS is responsible for the design and execution of the research, analysis of the data and drafting the paper.

## Supplemental Tables

**Supplemental Table 1.** Salaries for Predoctoral and Postdoctoral Trainees and Staff Scientists.

| NIH Guidelines |  | Other Source |  |
| --- | --- | --- | --- |
| Position | Salary | Position | Salary |
| Grad Student (NRSA) | \$25,320 | Grad Student (web average) | \$30,305 |
| Postdoc yr 1 (NRSA) | \$52,704 | Postdoc yr 1 (NSF SED) | \$48,000 (FY18) |
| Postdoc yr 5 (NRSA) | \$59,580 | | |
| Postdoc yr 7 (NRSA) | \$64,008 | | |
| Staff Scientist – GS13 step 1 | \$78,681 | Instructor (AAMC - median) | \$60,000 |
| Staff Scientist – GS14 step 1 | \$92,977 | Instructor (AAMC - 75<sup>th</sup> percentile) | \$72,000 |
| Staff Scientist – GS15 step 1 | \$109,366 | | |

**Supplemental Table 2.**
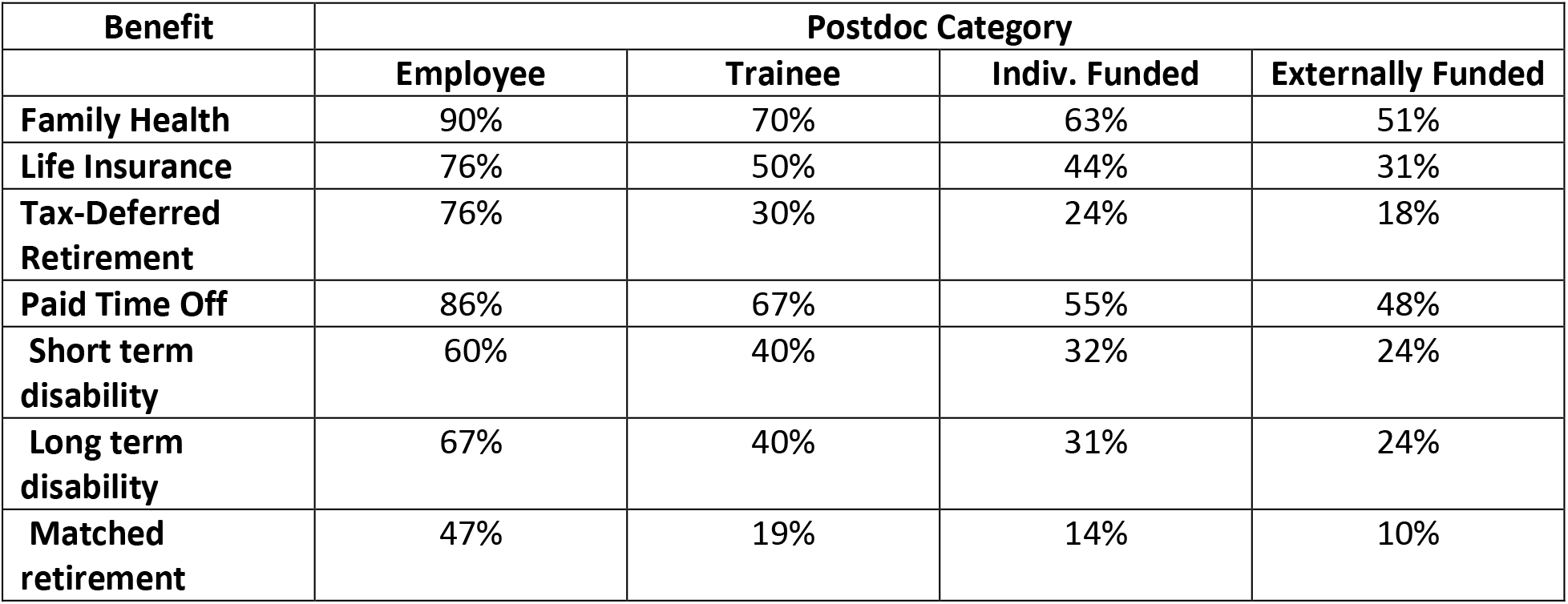
Postdoc Benefits – Percentage of Institutions Providing the Benefit.

**Supplemental Table 3.** Faculty Benefits at Institutions of Higher Education.

| Benefits | All combined |  | Public Institution |  | Private Institution |  |
| --- | --- | --- | --- | --- | --- | --- |
|  | % faculty covered | Value of benefit (% salary) | % faculty covered | Value of benefit (% salary) | % faculty covered | Value of benefit (% salary) |
| Retirement | 97.1 | 10.8 | 97.6 | 11.4 | 96.0 | 9.6 |
| Medical | 95.1 | 11.0 | 95.8 | 11.7 | 94.4 | 9.4 |
| **TOTAL** |  | 21.8 |  | 23.1 |  | 19.0 |
| Retirement excludes payments for unfunded retirement liability, prepaid retiree health insurance and social security. Medical excludes payments for long-term disability, Medicare and life insurance. |  |  |  |  |  |  |

**Supplemental Table 4.**
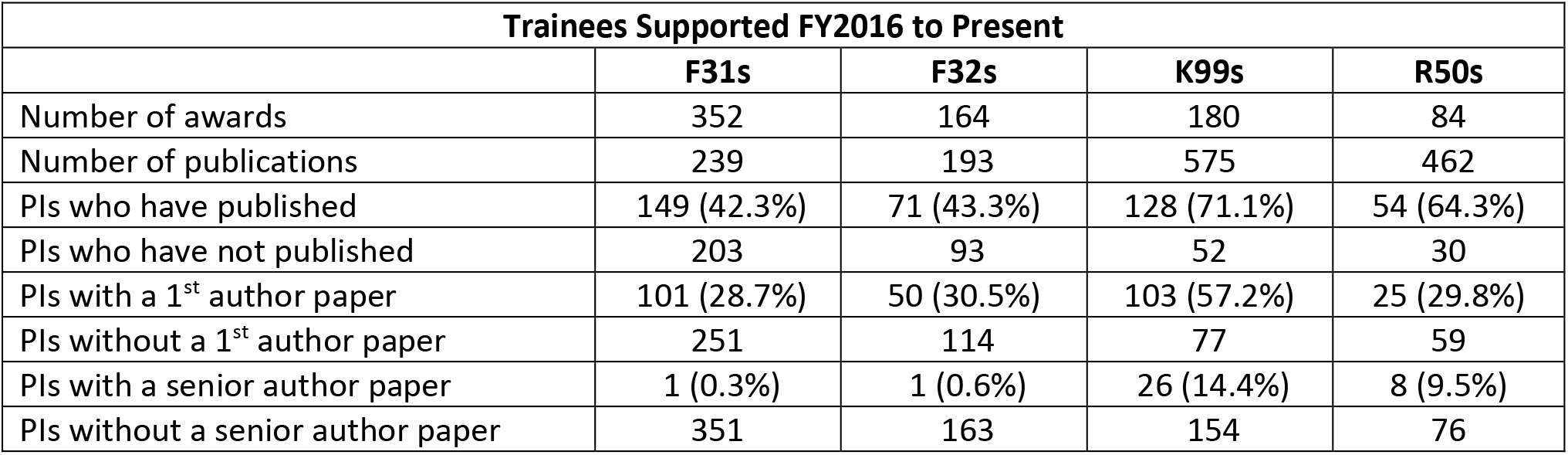
Profile of Publication Record of Trainees and Staff Scientists.

| **Trainees Supported FY2016 to Present** |  |  |  |  |
| --- | --- | --- | --- | --- |
|  | **F31s** | **F32s** | **K99s** | **R50s** |
| Number of awards | 352 | 164 | 180 | 84 |
| Number of publications | 239 | 193 | 575 | 462 |
| PIs who have published | 149 (42.3%) | 71 (43.3%) | 128 (71.1%) | 54 (64.3%) |
| PIs who have not published | 203 | 93 | 52 | 30 |
| PIs with a 1<sup>st</sup> author paper | 101 (28.7%) | 50 (30.5%) | 103 (57.2%) | 25 (29.8%) |
| PIs without a 1<sup>st</sup> author paper | 251 | 114 | 77 | 59 |
| PIs with a senior author paper | 1 (0.3%) | 1 (0.6%) | 26 (14.4%) | 8 (9.5%) |
| PIs without a senior author paper | 351 | 163 | 154 | 76 |

**Supplemental Table 5.** Profile of Publication Record of Trainees (FY2012-2016)

| **Trainees Supported During FY2012 to FY2016** |  |  |  |
| --- | --- | --- | --- |
|  | **F31s** | **F32s** | **K99s** |
| Number of awards | 3,605 | 3,588 | 1,252 |
| Number of publications | 9,576 | 10,886 | 6,166 |
| PIs who have published | 2,976 (82.6%) | 2,983 (83.1%) | 1,162 (92.8%) |
| PIs who have not published | 629 | 605 | 90 |
| PIs with a 1<sup>st</sup> author paper | 2,719 (75.4%) | 2,638 (73.5%) | 1,046 (83.6%) |
| PIs without a 1<sup>st</sup> author paper | 886 | 950 | 206 |
| PIs with a senior author paper | 90 (2.5%) | 221 (6.2%) | 435 (34.7%) |
| PIs without a senior author paper | 3,515 | 3,367 | 817 |
| Total citations of 1<sup>st</sup> author papers | 63,198 | 94,765 | 48,054 |
| Total citations of senior author papers | 2,087 | 4,358 | 7,495 |

